# OCTOPUS: Disk-based, Multiplatform, Mobile-friendly Metagenomics Classifier

**DOI:** 10.1101/2024.03.15.585215

**Authors:** Simone Marini, Alexander Barquero, Anisha Ashok Wadhwani, Jiang Bian, Jaime Ruiz, Christina Boucher, Mattia Prosperi

## Abstract

Portable genomic sequencers such as Oxford Nanopore’s MinION enable real-time applications in clinical and environmental health. However, there is a bottleneck in the downstream analytics when bioinformatics pipelines are unavailable, e.g., when cloud processing is unreachable due to absence of Internet connection, or only low-end computing devices can be carried on site. Here we present a platform-friendly software for portable metagenomic analysis of Nanopore data, the Oligomer-based Classifier of Taxonomic Operational and Pan-genome Units via Singletons (OCTOPUS). OCTOPUS is written in Java, reimplements several features of the popular Kraken2 and KrakenUniq software, with original components for improving metagenomics classification on incomplete/sampled reference databases, making it ideal for running on smartphones or tablets. OCTOPUS obtains sensitivity and precision comparable to Kraken2, while dramatically decreasing (4-to 16-fold) the false positive rate, and yielding high correlation on real-word data. OCTOPUS is available along with customized databases at https://github.com/DataIntellSystLab/OCTOPUS and https://github.com/Ruiz-HCI-Lab/OctopusMobile.

## Introduction

In the rapidly evolving field of microbial genomics, the ability to classify and analyze metagenomic data is crucial in both clinical and environmental health^1-4^. The advent of portable sequencing technologies, such as Oxford Nanopore’s MinION, has enabled real-time, in-field/point-of-care sequencing and analysis^5^. MinION’s outputs are long read nucleotide sequences, up to 4 million bases long, for a total throughput of ∼50 gigabytes. The device is powered by a laptop, weights only 87g, and it is very compact (105×23×37mm). There is also a tablet-incorporated version, the Mk1C (450g, 140×130×116mm), with preinstalled analytics software.

Several metagenomics classification tools are nowadays available for analyzing both short read data, e.g., Illumina, and long read data, e.g., PacBio and the aforementioned Nanopore^6^. The approaches can be roughly divided into: (i) database alignment, where the reads are aligned against a curated database of reference genomes; (ii) *k*-mer classifiers, which use lookup tables associating short sequences of fixed length k to taxa; and (iii) marker-based methods, where only species-specific gene families are considered. Group (i) includes traditional nucleotide/protein alignment programs such as BLAST, accelerated seed-chain aligners, e.g., DIAMOND^7^, Minimap2^8^, and aligners based on the Burrows-Wheeler transform, e.g., Centrifuge^9^. They usually provide the best sensitivity and specificity, low RAM usage, but are also the slowest. Burrows-Wheeler aligners provide increased speed over traditional ones but require more resources. The tools of group (ii) use different solutions for efficient *k*-mer hashing and linkage with taxa, can use taxonomy trees to classify at species, genus, or higher levels, and can work with both nucleotide and protein sequences. These methods are usually very fast, but hungry on RAM, and might have lower specificity than alignment-based methods. Popular tools include Clark Clark-S, Kaiju, Kraken2, and KrakenUniq^10-14^. The tools in the last group (iii) consider compact, curated, highly discriminative single marker gene databases, such as 16S, and combination of gene families for different microbial kingdoms, e.g., MetaPhlAn2^15^. In-depth reviews of the pros and cons of each approach, and comparisons on benchmark data have been made in multiple studies^16,17^.

Despite the widely availability of software and high classification accuracy –although it varies sensibly with reference databases and experimental data– the computational requirements for metagenomics classification often exceed the capabilities of regular desktops/laptops and by far those of mobile devices, creating a bottleneck in the process. The two major drawbacks of current metagenomics classifiers are the inability to be compiled and run on operating systems other than UNIX/Linux and the large amount of RAM needed. For instance, the Kraken2 documentation warns that “MacOS and other non-Linux operating systems are not explicitly supported by the developers” and it is noted that the MacOS does not support OpenMP, effectively limiting Kraken2 to run on a single thread. However, it is often possible to tweak the code and compile it on different platforms, e.g., Kraken and Minimap2 were shown to compile and run successfully on an Android device^18^, and we were able to do the same with Centrifuge on Windows using the developers’ suggestions. Regarding large RAM needs, some tools like Kraken2 and KrakenUniq allow to swap on disk, although the database indexing still requires large RAM allocation. Minor issues include the complexity of setting up the software pipelines, from installing the right dependencies, to code compilation and command scripts with optimized parameters and up-to-date databases. Most minor issues can be solved with standardized analytic protocols^19^ and proper IT support, but the problem becomes non-trivial in mobile settings or remote locations where internet connection, IT personnel, and other support is not readily available. For these reason, all-in-one solutions have been proposed, such as the Mk1C, and open-source mobile apps, e.g., iGenomics for sequence alignment^20^ and KARGAMobile for classification of antimicrobial resistance^21^.

Inspired by Kraken2 and KrakenUniq, we here present a novel product for platform-friendly, portable metagenomics analysis of long read data, the Oligomer-based Classifier of Taxonomic Operational and Pan-genome Units via Singletons (OCTOPUS). Our tool harnesses the methodological strengths of its predecessors, with a number of novelties, including a statistical criterion for read classification that dramatically decreases the false positive rate without increasing the false negative rate. OCTOPUS eliminates the need for high-performance computing resources, simplifies dependencies and installation on to multiple operating systems –the code is fully written in Java– allowing usage directly on tablets or smartphones connected to the MinION sequencer.

## Methods

As mentioned, OCTOPUS reimplements several algorithmic and data structure components from Kraken2 and KrakenUniq. Like them, OCTOPUS stores *k*-mer minimizers^22^ instead of *k*-mers, and uses the HyperLogLogPlus^23,24^ algorithm for keeping counts of distinct *k*-mer minimizers found for each species.

Nonetheless, there are also some notable differences and innovations, namely: (i) *k*-mer minimizer representation and handling; (ii) clustering of reference genomes and *k*-mer filtering; (iii) compact database indexing; (iv) probabilistic classification criterion for the single read and summary of species’ coverage and depth for a read sample.

### Representation and handling of k-mers

The *k*-mer representation in OCTOPUS uses a polynomial rolling hash^25^. Given a constant a, and constant integer values for input characters *c*_*1*_, *… c*_*m*_, the hash is defined as *H = c*_*1*_*a*^*k-1*^ *+ c*_*2*_*a*^*k2*^ *+ … + c*_*k*_*a*^*0*^. The hash calculations are done modulo *n* (usually a large prime). For nucleotide sequences, composed by the 4 bases A, C, G, T, we can set the characters *c*_*1*_, *… c*_*4*_ as 0 (i.e., A) to 4 (i.e., T), and the constant a to 4. When *k* is fixed, the hash values respect alphabetical order. If a sequence includes ambiguous bases or erroneous characters, an additional masking character, N, can be added, increasing *a* to 5. With 64-bit integers, the maximum *k*-mer length that can be hashed without modulo overflow, i.e., no collisions, is 31 (27 using masking). OCTOPUS uses read minimizers in the same way as Kraken2 does, with a *l*-nucleotide difference between the minimizer and the *k*-mer. We set *l* to 4, thus the maximum *k*-mer length becomes 35 (31 using masking). In the rolling hash, removal of a heading character or addition of a trailing character involves only subtraction or addition, while shifting involves division by a constant (or multiplication by its inverse). These properties allow the calculation of all *k*-mers in a read by single character removal, shift, and addition consecutively. To make *k*-mer calculation efficient, we use a precomputed matrix for all *c*_*j*_*a*^*k-j*^ values and only subtraction and multiplication for *k*-mer shifting. Additionally, we employ a fixed-size character array for the read, and two independent hash tables that map characters to integers, one for forward-strand and another for reverse complement sequences (with complementary integers); in this way, we scan the character array only twice, from beginning to end and vice versa. When comparing *k*-mers and their reverse complements, we retain the one with the lowest alphabetical order, i.e., the smallest integer hash. The minimizers are calculated efficiently, since we compute all *(k-l)*-mers and then update the *k*-mer minimizer over a sliding window of length *l*. Currently, OCTOPUS does not use spaced seeds. **Figure 1A** illustrates the *k*-mer minimizer set calculation on a single read example (*k*=7, *l*=2), omitting reverse complements for ease of reading.

**Figure 1.**
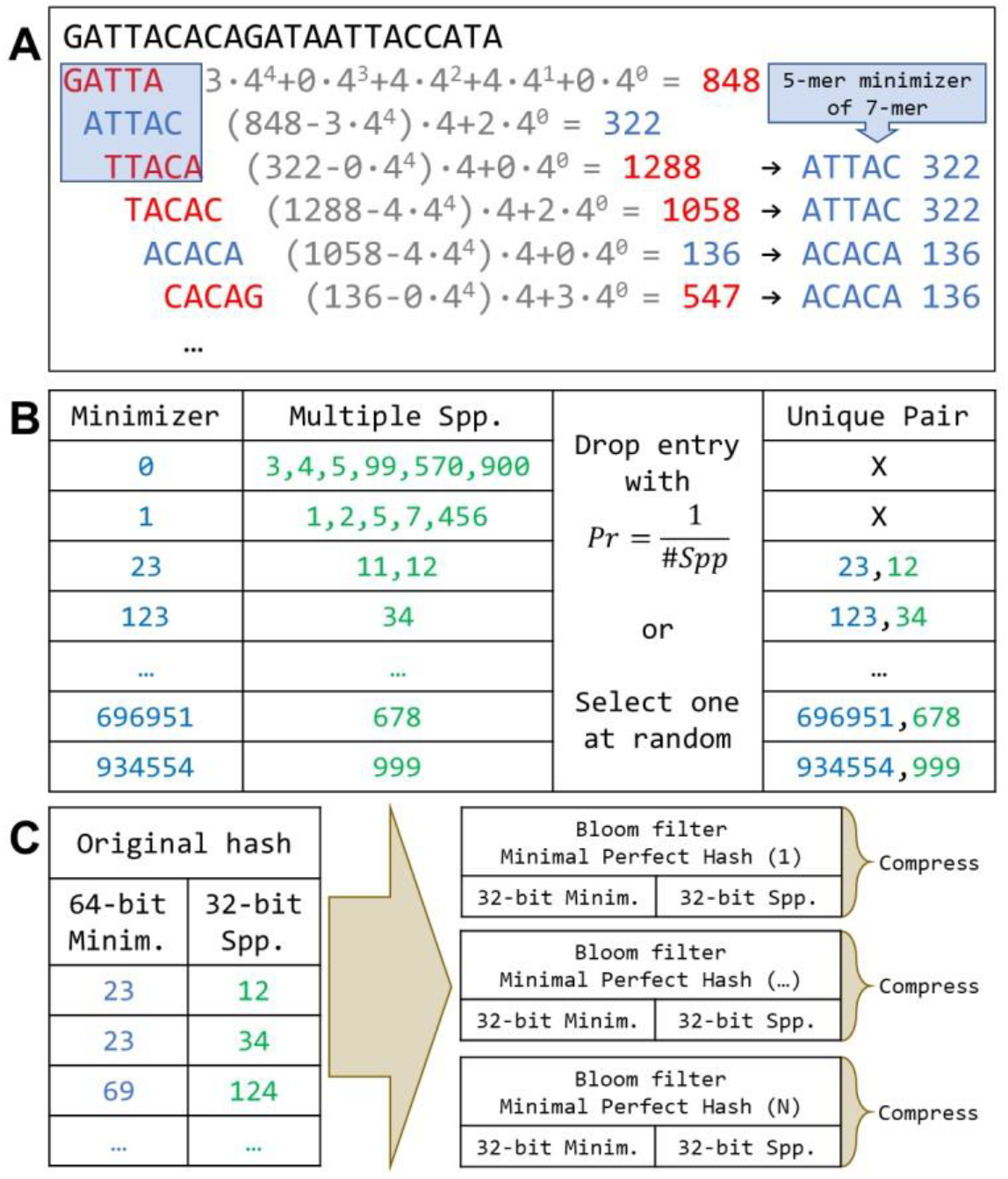
Calculation of *k*-mer minimizers using rolling hash (A), *k*-mer probabilistic filtering based on species’ cooccurrence (B), and compact (64-to 32-bit) *k*-mer indexing through Bloom filters, minimal perfect hashing and compression (C).

### Species’ clustering and k-mer filtering

Kraken2 uses the National Center for Biotechnology Information (NCBI) GenBank taxonomy tree to output classifications at the species, genus or higher levels based on the lowest common ancestor algorithm^26^. However, this approach presents a major drawback: When a species’ reference database contains several organisms with high intra-species similarity, Kraken2 often cannot decide on the species and reports results at the genus level^27^. OCTOPUS does not use the taxonomy tree; rather it uses an approach that relies on (i) species/genomes’ clustering and (ii) exploitation of *k*-mer uniqueness among species. First, OCTOPUS runs a leader clustering^28^ of all species based on a modified Jaccard similarity, i.e., the number of *k*-mer minimizers in common divided by the minimum of the two respective totals, that is, |A∩B| / min(|A|,|B|). The threshold similarity for the leader clustering is an input parameter set to 90% as default value. In this way, OCTOPUS can classify a read either as a single species, or as part of a cluster of species. Second, OCTOPUS uses a probabilistic filtering of *k*-mer minimizers that are shared by multiple species. If minimizer is shared by *N* species, it is filtered out with a probability of 1/*N*. If not dropped, it is assigned randomly to one of the *N* species (**Figure 1B**). This probabilistic filtering has two advantages: (i) it naturally eliminates low-complexity *k*-mer minimizers, and (ii) it creates a one-to-one relationship between *k*-mer minimizers and species.

### Compact database indexing

For *k*-mer storage and querying, OCTOPUS uses a novel approach for the compact hash table that links *k*-mer minimizers to species. We devise a probabilistic minimal perfect hashing algorithm that combines a Bloom filter on top of a regular hashing and compression, reducing the space needed for *k*-mer minimizers of more than a half. In detail, each 64-bit *k*-mer minimizer is added to a Bloom filter and at the same time hashed into a 32-bit integer, until a collision is found; colliding items are removed and stored for later use. Once all minimizers have been processed, the process restarts with a new Bloom filter and a new hash table on the leftovers, and so on until all minimizers have been bucketed. This algorithm improves that of Limasset et al.^29^, which cannot resolve queries when an item is not part of the key set, while ours can. At the end of the procedure, all tables are compressed using Delta/LZF or Deflate/LZF for the mobile version. Before compression, the number of bits for one minimizer reduces from 64 to 32, plus the overhead of the Bloom filter and of the disk-based hash structures. **Figure 1C** summarizes the steps for the creation of this set of compressed minimal perfect hash tables.

### Probabilistic read classification and sample summaries

The classification criterion of a single read in OCTOPUS is based on the maximum number of unique *k*-mer minimizers that are found among species or clusters of species, and such number must exceed a given threshold minimizing the probability of being a false positive. The need to add a threshold becomes important with long read data, since the probability that a *k*-mer minimizer is found by chance in a given genome increases with the read length. Also, it has been shown that customized, taxon-specific reference databases increase the probability of false positives^30^. We previously provided both theoretical and empirical determination of distributions of *k*-mer frequencies over genomes under various assumptions, e.g., Markovian^31,32^. However, OCTOPUS uses *k*-mer minimizers instead of *k*-mers, and thus a direct formula is more complicated to obtain, even under a simplified zero-order Markov and Poisson distribution assumption. In fact, estimating the density of minimizers is not trivial. In the work by Roberts et al.^22^ that introduced minimizers, the proportion of *k*-mer minimizers in a sequence of length n was estimated to be roughly equal to 2/(*w*+1), where *w* is the window length, i.e., *k-(k-l)*+1 = *l*+1. Refined estimates of the lower and upper bounds, including optimal theoretical densities for given minimizers’ schemes, have been derived^33,34^. For OCTOPUS, we use an empirical estimate, tuned on the reference database that is indexed and used for classification. In detail, we generate up to 1,000 random reads (with equal nucleotide probability) with read lengths ranging from *k-l* to 8 million base pairs (incremented by √√2) and we calculate the average maximum number of *k*-mer minimizer hits to a single species. We assume that the number of hits follows a Poisson distribution, thus we can calculate the probability that t matches to a species happen by chance, and use Pr(*t*) as a customizable threshold.

The summary of species’ depth and coverage is done as follows: each read assigned to a species or to a species’ cluster increases to the depth count by one unit, and all the distinct *k*-mer minimizers of that read are added to the species’ HyperLogLogPlus counter if they match the species’ *k*-mer spectrum. The species’ coverage is calculated as the ratio between the final HyperLogLogPlus count after processing all reads and the actual number of distinct *k*-mer minimizers.

### Application implementation

OCTOPUS is written in Java, using the standard v.8 core library plus greplin-bloomfilter, stream-lib-3.0, mapdb-3.0, and h2-mvstore-2.2 (for the mobile version) with respective dependencies. The genome references’ indexing modules can be run with very limited hardware resources, as little as a few hundred megabytes of RAM, since the programs will automatically swap on to disk if needed; there are no major drawbacks, except that the number of files and size of the final minimal perfect hashes might be larger, besides longer runtimes. The user can limit the final reference database size to a desired number of gigabytes (a portion of the *k*-mer minimizers will be discarded at random). The hash tables are stored into sorted memory mapped files with binary search key retrieval. For the mobile version, b-trees are used, because the Java library for memory mapped files (mapdb-3.0) contains unsafe code. The classifier works exclusively on-disk; thus, the RAM usage is minimal as well.

### Android porting

We adapt the OCTOPUS code to run natively as a console application on the Android platform. The application is written in Java v.21 and developed using Android Studio IDE (version Hedgehog | 2023.1.1 Patch 2), targeting API level 31 with a minimum supported API level of 28. Due to existing Android constraints regarding execution time on regular worker threads, which have a ten-minute limit, we employ high-priority foreground asynchronous threads with long-running capabilities. This approach allows for extended analysis durations without interruptions, which could happen depending on the characteristics and size of the database or the reads files. It also provides the functionality required to analyze the application performance simultaneously. To address Android’s file access restrictions, especially concerning third-party libraries, some minor modifications are made, compared to the desktop version of OCTOPUS. In particular, the third-party library greplin-bloom-filter-master has methods that require accessing files directly. However, Android restricts third-party library file access to the exclusive space of the internal storage. To circumvent this limitation, the database and reads files are included within the application installation package and subsequently stored in the app’s internal storage. This ensures compliance with Android’s security protocols while allowing the code to function with minimal changes to the desktop version.

All code is available on GitHub (https://github.com/DataIntellSystLab/OCTOPUS for the general-purpose version and https://github.com/Ruiz-HCI-Lab/OctopusMobile for the Android porting) and licensed under Apache 2.0.

### Evaluation framework

The experimental testbed for OCTOPUS is threefold: (i) we evaluate its performance in classifying simulated bacterial (true positives) and non-bacterial (true negatives) long Nanopore reads on a real, minimally redundant bacterial reference database, comparing it with Kraken2; (ii) we evaluate its capability to run on an Android smartphone, using real Nanopore data against customized databases (viral and bacterial); (iii) we evaluate its agreement with Kraken2 over real-world metagenomics samples. It is important to note that the comparison with Kraken2, from classification performance to time of execution, is made by indexing Kraken2 over the very same databases.

To create the minimally redundant bacterial reference database, we consider all the available complete and representative bacterial genomes of the Bacterial and Viral Bioinformatics Resource Center (BV-BRC) v.3.31.1235, using the NCBI taxonomy to label each genome at species level. Both OCTOPUS and Kraken2 databases are indexed on these genomes.

The bacterial (positive) test set is made as follows. For each genome in the reference database, we identify and download a different, random genome corresponding to the specific taxon reported in BV-BRC, using the NCBI datasets tool (https://github.com/ncbi/datasets). We use each FASTA file of said genomes to simulate Nanopore reads (FASTQ files) via the PBSIM2 tool36 with default parameters (minimum length = 100; maximum length = 1,000,000; mean length = 9,000; length standard deviation = 7,000; minimum accuracy = 0.75, maximum accuracy = 1, mean accuracy = 0.85) and the recommended Nanopore difference-ratio = 23:31:46. We set depth = 5. True positives (TP) and false positives (FP) are calculated read-by-read for each simulated genome, at the species level for both Kraken2 and OCTOPUS (except for a species’ cluster, in which case the exact species must be contained in that cluster). False negatives (FN) are counted as the total of the reads for the target species, minus the sum of TP and FP.

The non-bacterial (negative) test sets include two datasets, one made with mammalian genomes and the other with viral genomes. The mammalian test set includes all the available genomes in NCBI RefSeq (as of Jan/06/2024) pertaining to the Camelidae family (NCBI taxon ID 9835). We concatenate the FASTA files, and extract 5,000 random sequences, which are used to generate the reads via PBSIM2 as explained above. The viral test set includes available reference genomes (as of Dec/18/2023). Reads are simulated via PBSIM2 as well, but for the viral test set we set depth = 25. On a whole dataset, we count any matched read, i.e., different from unclassified, as a false positive (FP), and any unmatched read as a true negative (TN).

The Android customized databases include: (i) all viral genomes from RefSeq (as of Mar/5/2023); (ii) bacterial genomes from the BV-BRC reference set, selecting all species (and up to three species per genus) of concern for antibiotic resistance as determined by the World Health Organization (https://www.who.int/news/item/27-02-2017who-publishes-list-of-bacteria-for-which-new-antibiotics-are-urgently-needed); (iii) the MEGARes v.3.0 database, contains sequence data for nearly 9,000 hand-curated antimicrobial resistance genes37. The test sets included Nanopore metagenomics sequencing data generated from biological samples of hospitalized individuals with severe pneumonia38.

The comparison of OCTOPUS and Kraken2 over real-world data is done using data from the MetaSUB consortium^39^. MetaSUB is an international project collecting and sequencing surface samples from public transportation systems worldwide, with the goal of characterizing urban microbiomes. We select at random 100 MetaSUB metagenomics experiments, carried out using Illumina sequencing technology (short reads), with the corresponding sequencing (paired) FASTQ files. We apply quality control as described in our previous work^40^ and capped each FASTQ pair to ∼50MB, for a total of ∼5GB (compressed) data. Then Kraken2 and OCTOPUS are run over each capped sample with parameters and bacterial genome references described above.

For replication purposes, all the code scripts used to generate the data sets, the NCBI taxon IDs for each reference database and test are available on the Github repository links provided.

## Results

The initial bacterial reference database downloaded from BV-BRC includes 4,234 genomes, corresponding to 4,188 unique species according to the NCBI taxonomy. We generate ∼10 million long Nanopore reads for the (positive) bacterial test set, totaling 75 GB (compressed). Of note, 30 of the initial FASTA files (less than 1%) are discarded because too short or not available at the assembly level, for a total of 4,202 retained genomes. The mammalian test set includes 7 genomes (241,673 reads, 1.7 GB, compressed), while the viral test set includes 47 genomes (3,621 reads, 18 MB, compressed).

The performance distribution summaries for all test sets are illustrated in Figure 2. As explained in the methods, the TP rate (sensitivity) and precision (sensitivity × prevalence) for the positive test set are calculated per-species, whilst the FP rate is based on whole negative sets. In detail, on the bacterial (positive) test set, OCTOPUS yields a median (interquartile range, IQR) sensitivity of 0.944 (0.896-0.952), and a precision of 0.999 (0.982-1), whilst Kraken2 yields a sensitivity of 0.945 (0.901-0.95) and a precision of 0.991 (0.968-0.996). On the mammalian (negative) test set, the mean (± standard error) FP rate is 0.007 (±0.0002) for OCTOPUS and 0.112 (±0.0006) for Kraken2. On the viral (negative) test set, it is 0.021 (±0.002) for OCTOPUS and 0.082 (±0.005) for Kraken2. Overall, OCTOPUS decreases the FP rate between 4 and 16 folds.

**Figure 2.**
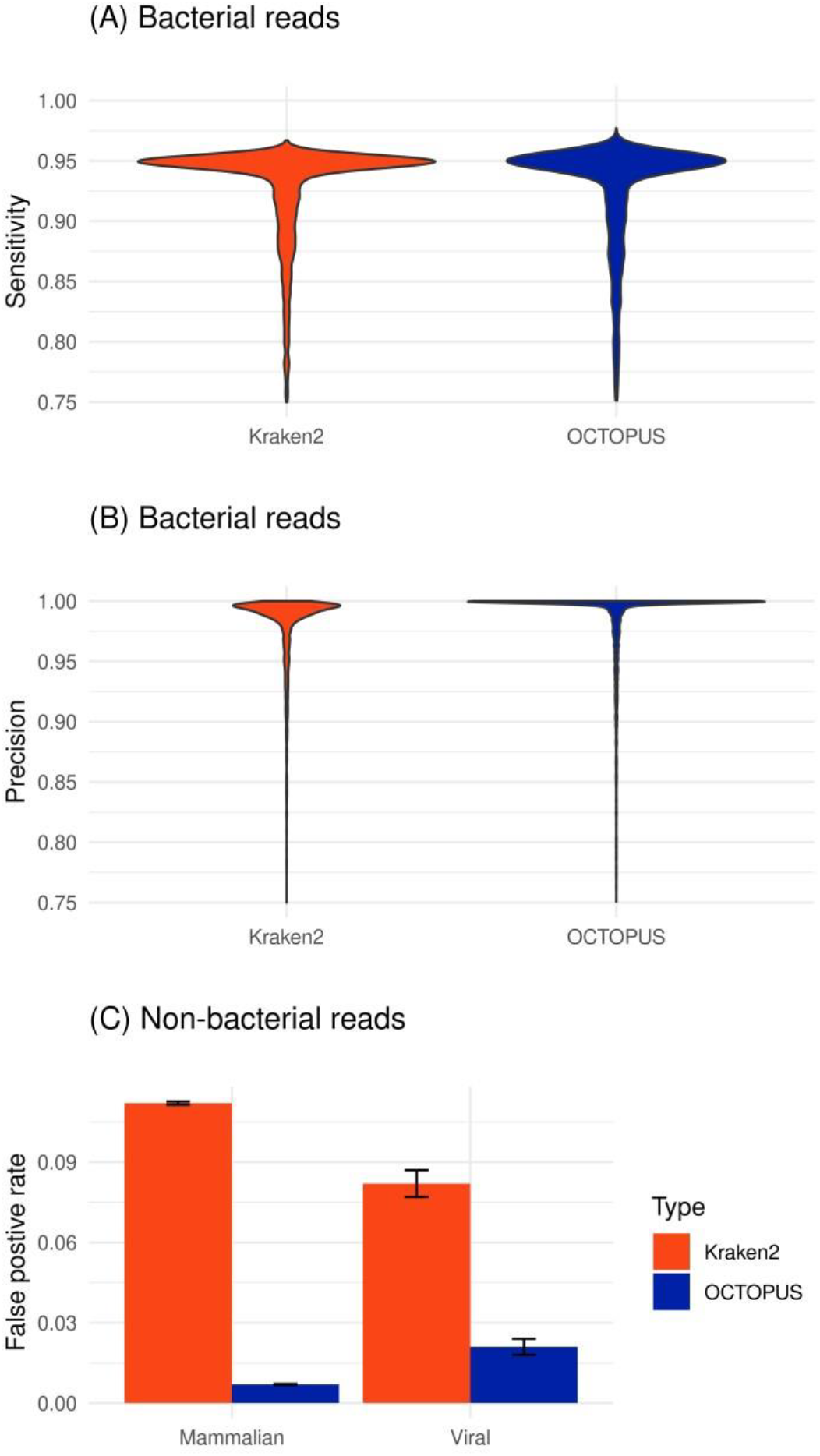
Comparison of classification performance between OCTOPUS and Kraken2. Panel (A) and (B) show the distribution of sensitivity and precision on simulated bacterial datasets (4,202 genomes, ∼10 million reads), while panel (C) shows the false positive rate on non-bacterial datasets made of mammalian and viral genomes (7 genomes and 241,673 reads, 47 genomes and 3,621 reads, respectively).

Sensitivity and precision are strongly correlated on the (positive) bacterial test set, 0.92 and 0.93 respectively (Pearson). However, if we limit the analysis to the distribution tails with lowest scores, the correlation drops. By considering the union of bacterial genomes with bottom 5% (i.e., 210 genomes) performances for each algorithm, we obtain a 0.45 correlation (Pearson) for precision (total genomes: 226); and a 0.39 correlation (Pearson) for sensitivity (total genomes: 237).

In terms of speed, on a machine with 16 AMD EPYC 7702 CPUs and 8GB RAM, OCTOPUS is on average 1.6x slower than Kraken2, using the memory mapped option to process data exclusively on the disk, on the test sets. The runtime and memory usage of the Android porting are given in Table 1. Of note, when trying to expand the bacterial database by including more species per genus (up to a total of 400+), the Android app crashes or runs out of memory when the b-tree store grows larger than 3GB. The average and maximum temperatures are high, but the device does not shut off due to overheating.

**Table 1.** Runtime and memory usage of OCTOPUS’ Android porting on real metagenomics datasets (clinical samples from hospitalized individuals with pneumonia).

| Database type | OCTOPUS database size | FASTQ Accession ID | FASTQ Size / Gzip (MB) | CPU / Wall time (minutes) | Average / Max RAM (MB) | Average / Max Temp (C°) |
| --- | --- | --- | --- | --- | --- | --- |
| Viral RefSeq | 14,517 species/clusters<br>97M *k*-mers<br>909 MB | SRX6447019 | 847 / 138 | 7 / 9 | 363 / 442 | 56 / 70 |
|  |  | SRX6447061 | 1050 / 172 | 8 / 12 | 365 / 452 | 54 / 66 |
|  |  | SRX6447018 | 1740 / 290 | 16 / 21 | 358 / 441 | 53 / 66 |
| Bacterial (WHO concern) | 111 species/clusters<br>121M *k*-mers<br>984 MB | SRX6447019 | 847 / 138 | 16 / 18 | 206 / 265 | 55 / 72 |
|  |  | SRX6447061 | 1050 / 172 | 16 / 19 | 189 / 236 | 54 / 70 |
|  |  | SRX6447018 | 1740 / 290 | 40 / 42 | 198 / 259 | 53 / 71 |
| MEGARes | 3,990 gene/clusters<br>1M *k*-mers<br>10 MB | SRX6447019 | 847 / 138 | 3 / 6 | 135 / 183 | 55 / 66 |
|  |  | SRX6447061 | 1050 / 172 | 4 / 8 | 125 / 178 | 54 / 65 |
|  |  | SRX6447018 | 1740 / 290 | 8 / 13 | 132 / 185 | 54 / 64 |

For the comparison over real-world metagenomics data, i.e., MetaSUB, we compare OCTOPUS and Karaken2 classification outputs by measuring the per-species pairwise correlation of mapped reads for each sample. The results indicate a strong agreement between the two approaches, with median (IQR) Pearson correlation of 0.99 (0.98, 1), and median (IQR) Spearman correlation of 0.87 (0.78, 0.9).

## Discussion and Conclusions

OCTOPUS demonstrates robust classification performance with long read Nanaopore data and the bacterial reference database, effectively sorting out non-bacterial reads. OCTOPUS and Kraken2 have practically indistinguishable sensitivity and precision, but OCTOPUS dramatically decreases the false negative rates (4-to 16-fold better than Kraken2). The Android app runs smoothly on the customized databases, although the implementation reaches a practical limit when the index surpasses 3GB. In any case, *k*-mer subsampling is available like in Kraken2, and multiple databases can be combined thanks to the robustness in false negative rates. Furthermore, when tested over real-world Illumina reads, OCTOPUS shows strong species classification correlation with Kraken2.

This work has some limitations. OCTOPUS’ Java code runs sensibly slower than Kraken2, whose core is written in C++. The advantage of OCTOPUS is that it works on any device that mounts a Java virtual machine, but there are some drawbacks regarding the optimization of data structures, code, hardware acceleration, and speed of execution. In terms of data structures, we recognize that further improvements could be done by reducing the bits allocated to represent species, and that a customized memory-mapped approach to the hash tables could be used instead of the off-the-shelf mapdb. Also, the data processing is exclusively on-disk, while a solution that loads chunks of hashes on RAM could be devised. Further, the code for *k*-mer extraction, filtering, genome clustering and database indexing is not optimized, resulting in very slow preprocessing times. From an algorithmic standpoint, one limitation of OCTOPUS is that spaced seeds are not implemented, even though it has been shown that they can improve classification accuracy^40^. One reason is that spaced seeds increase the complexity of *k*-mer calculation, especially for the rolling hash; however, some solutions could be adapted to our approach^41^. Another limitation related to the algorithm is that the genome clustering is not based on species’ taxonomy and that the leader clustering is order dependent. Finally, it would be interesting to derive an exact formula for the expected number of hits from random sequences, rather than rely on an empirical approximation.

In conclusion, the OCTOPUS software is ideal for mobile applications using long read data and ad hoc databases, where computing resources are limited, and it is crucial to minimize false positives.

## Acknowledgments

This work was in part supported by US federal grants: NIH NIAID R01AI145552; NIH NIAID R01 AI170187; NIAID 1R01AI141810; NSF SCH 2013998.

## References

1. C. Staley and M. J. Sadowsky, “Practical considerations for sampling and data analysis in contemporary metagenomics-based environmental studies,” J. Microbiol. Methods, vol. 154, pp. 14–18, Nov. 2018, doi: 10.1016/j.mimet.2018.09.020.

2. R. Knight et al., “Best practices for analysing microbiomes,” Nat. Rev. Microbiol., vol. 16, no. 7, pp. 410–422, Jul. 2018, doi: 10.1038/s41579-018-0029-9.

3. C. Y. Chiu and S. A. Miller, “Clinical metagenomics,” Nat. Rev. Genet., vol. 20, no. 6, pp. 341–355, Jun. 2019, doi: 10.1038/s41576-019-0113-7.

4. N. Li, Q. Cai, Q. Miao, Z. Song, Y. Fang, and B. Hu, “High-Throughput Metagenomics for Identification of Pathogens in the Clinical Settings,” Small Methods, vol. 5, no. 1, p. 2000792, Jan. 2021, doi: 10.1002/smtd.202000792.

5. J. Pugh, “The Current State of Nanopore Sequencing,” Methods Mol. Biol. Clifton NJ, vol. 2632, pp. 3–14, 2023, doi: 10.1007/978-1-0716-2996-3_1.

6. F. P. Breitwieser, J. Lu, and S. L. Salzberg, “A review of methods and databases for metagenomic classification and assembly,” Brief. Bioinform., vol. 20, no. 4, pp. 1125–1136, Jul. 2019, doi: 10.1093/bib/bbx120.

7. B. Buchfink, C. Xie, and D. H. Huson, “Fast and sensitive protein alignment using DIAMOND,” Nat. Methods, vol. 12, no. 1, Art. no. 1, Jan. 2015, doi: 10.1038/nmeth.3176.

8. H. Li, “Minimap2: pairwise alignment for nucleotide sequences,” Bioinformatics, vol. 34, no. 18, pp. 3094–3100, Sep. 2018, doi: 10.1093/bioinformatics/bty191.

9. D. Kim, L. Song, F. P. Breitwieser, and S. L. Salzberg, “Centrifuge: rapid and sensitive classification of metagenomic sequences,” Genome Res., vol. 26, no. 12, pp. 1721–1729, Dec. 2016, doi: 10.1101/gr.210641.116.

10. R. Ounit, S. Wanamaker, T. J. Close, and S. Lonardi, “CLARK: fast and accurate classification of metagenomic and genomic sequences using discriminative k-mers,” BMC Genomics, vol. 16, no. 1, p. 236, Mar. 2015, doi: 10.1186/s12864-015-1419-2.

11. R. Ounit and S. Lonardi, “Higher classification sensitivity of short metagenomic reads with CLARK-S,” Bioinforma. Oxf. Engl., vol. 32, no. 24, pp. 3823–3825, Dec. 2016, doi: 10.1093/bioinformatics/btw542.

12. P. Menzel, K. L. Ng, and A. Krogh, “Fast and sensitive taxonomic classification for metagenomics with Kaiju,” Nat. Commun., vol. 7, no. 1, Art. no. 1, Apr. 2016, doi: 10.1038/ncomms11257.

13. D. E. Wood, J. Lu, and B. Langmead, “Improved metagenomic analysis with Kraken 2,” Genome Biol., vol. 20, no. 1, p. 257, Nov. 2019, doi: 10.1186/s13059-019-1891-0.

14. F. P. Breitwieser, D. N. Baker, and S. L. Salzberg, “KrakenUniq: confident and fast metagenomics classification using unique k-mer counts,” Genome Biol., vol. 19, no. 1, p. 198, Nov. 2018, doi: 10.1186/s13059-018-1568-0.

15. D. T. Truong et al., “MetaPhlAn2 for enhanced metagenomic taxonomic profiling,” Nat. Methods, vol. 12, no. 10, Art. no. 10, Oct. 2015, doi: 10.1038/nmeth.3589.

16. S. H. Ye, K. J. Siddle, D. J. Park, and P. C. Sabeti, “Benchmarking Metagenomics Tools for Taxonomic Classification,” Cell, vol. 178, no. 4, pp. 779–794, Aug. 2019, doi: 10.1016/j.cell.2019.07.010.

17. J. Marić, K. Križanović, S. Riondet, N. Nagarajan, and M. Šikić, “Comparative analysis of metagenomic classifiers for long-read sequencing datasets,” BMC Bioinformatics, vol. 25, no. 1, p. 15, Jan. 2024, doi: 10.1186/s12859-024-05634-8.

18. M. Oliva, F. Milicchio, K. King, G. Benson, C. Boucher, and M. Prosperi, “Portable nanopore analytics: are we there yet?,” Bioinformatics, vol. 36, no. 16, pp. 4399–4405, Apr. 2020, doi: 10.1093/bioinformatics/btaa237.

19. J. Lu et al., “Metagenome analysis using the Kraken software suite,” Nat. Protoc., vol. 17, no. 12, pp. 2815–2839, Dec. 2022, doi: 10.1038/s41596-022-00738-y.

20. A. Palatnick, B. Zhou, E. Ghedin, and M. C. Schatz, “iGenomics: Comprehensive DNA sequence analysis on your Smartphone,” GigaScience, vol. 9, no. 12, p. giaa138, Dec. 2020, doi: 10.1093/gigascience/giaa138.

21. A. Barquero, S. Marini, C. Boucher, J. Ruiz, and M. Prosperi, “KARGAMobile: Android app for portable, realtime, easily interpretable analysis of antibiotic resistance genes via nanopore sequencing,” Front. Bioeng. Biotechnol., vol. 10, 2022, Accessed: Jan. 06, 2024. [Online]. Available: 10.3389/fbioe.2022.1016408

22. M. Roberts, W. Hayes, B. R. Hunt, S. M. Mount, and J. A. Yorke, “Reducing storage requirements for biological sequence comparison,” Bioinforma. Oxf. Engl., vol. 20, no. 18, pp. 3363–3369, Dec. 2004, doi: 10.1093/bioinformatics/bth408.

23. P. Flajolet, É. Fusy, O. Gandouet, and F. Meunier, “HyperLogLog: the analysis of a near-optimal cardinality estimation algorithm,” Discrete Math. Theor. Comput. Sci., vol. DMTCS Proceedings vol. AH,…, no. Proceedings, p. 3545, Jan. 2007, doi: 10.46298/dmtcs.3545.

24. S. Heule, M. Nunkesser, and A. Hall, “HyperLogLog in practice: algorithmic engineering of a state of the art cardinality estimation algorithm,” in Proceedings of the 16th International Conference on Extending Database Technology, Genoa Italy: ACM, Mar. 2013, pp. 683–692. doi: 10.1145/2452376.2452456.

25. R. M. Karp and M. O. Rabin, “Efficient randomized pattern-matching algorithms,” IBM J. Res. Dev., vol. 31, no. 2, pp. 249–260, Mar. 1987, doi: 10.1147/rd.312.0249.

26. D. H. Huson, A. F. Auch, J. Qi, and S. C. Schuster, “MEGAN analysis of metagenomic data,” Genome Res., vol. 17, no. 3, pp. 377–386, Mar. 2007, doi: 10.1101/gr.5969107.

27. D. J. Nasko, S. Koren, A. M. Phillippy, and T. J. Treangen, “RefSeq database growth influences the accuracy of k-mer-based lowest common ancestor species identification,” Genome Biol., vol. 19, no. 1, p. 165, Oct. 2018, doi: 10.1186/s13059-018-1554-6.

28. J. A. Hartigan, Clustering Algorithms, 99th ed. USA: John Wiley & Sons, Inc., 1975.

29. A. Limasset, G. Rizk, R. Chikhi, and P. Peterlongo, “Fast and scalable minimal perfect hashing for massive key sets.” arXiv, Feb. 16, 2017. doi: 10.48550/arXiv.1702.03154.

30. V. R. Marcelino, E. C. Holmes, and T. C. Sorrell, “The use of taxon-specific reference databases compromises metagenomic classification,” BMC Genomics, vol. 21, no. 1, p. 184, Feb. 2020, doi: 10.1186/s12864-020-6592-2.

31. M. C. F. Prosperi, L. Prosperi, R. R. Gray, and M. Salemi, “On counting the frequency distribution of string motifs in molecular sequences,” Int. J. Biomath., vol. 05, no. 06, p. 1250055, Nov. 2012, doi: 10.1142/S1793524512500556.

32. M. Prosperi, S. Marini, and C. Boucher, “Fast and exact quantification of motif occurrences in biological sequences,” BMC Bioinformatics, vol. 22, no. 1, p. 445, Sep. 2021, doi: 10.1186/s12859-021-04355-6.

33. G. Marçais, D. DeBlasio, and C. Kingsford, “Asymptotically optimal minimizers schemes,” Bioinforma. Oxf. Engl., vol. 34, no. 13, pp. i13–i22, Jul. 2018, doi: 10.1093/bioinformatics/bty258.

34. H. Zheng, C. Kingsford, and G. Marçais, “Improved design and analysis of practical minimizers,” Bioinforma. Oxf. Engl., vol. 36, no. Suppl_1, pp. i119–i127, Jul. 2020, doi: 10.1093/bioinformatics/btaa472.

35. R. D. Olson et al., “Introducing the Bacterial and Viral Bioinformatics Resource Center (BV-BRC): a resource combining PATRIC, IRD and ViPR,” Nucleic Acids Res., vol. 51, no. D1, pp. D678–D689, Jan. 2023, doi: 10.1093/nar/gkac1003.

36. Y. Ono, K. Asai, and M. Hamada, “PBSIM2: a simulator for long-read sequencers with a novel generative model of quality scores,” Bioinforma. Oxf. Engl., vol. 37, no. 5, pp. 589–595, May 2021, doi: 10.1093/bioinformatics/btaa835.

37. N. Bonin et al., “MEGARes and AMR++, v3.0: an updated comprehensive database of antimicrobial resistance determinants and an improved software pipeline for classification using high-throughput sequencing,” Nucleic Acids Res., vol. 51, no. D1, pp. D744–D752, Jan. 2023, doi: 10.1093/nar/gkac1047.

38. L. Yang et al., “Metagenomic identification of severe pneumonia pathogens in mechanically-ventilated patients: a feasibility and clinical validity study,” Respir. Res., vol. 20, no. 1, p. 265, Nov. 2019, doi: 10.1186/s12931-0191218-4.

39. Mason C., et al., “The metagenomics and Metadesign of the subways and urban biomes (MetaSUB) international consortium inaugural meeting report,” Microbiome vol. 4 no. 24, Jun 2016, doi: 10.1186/s40168-016-0168-z.

40. Marini S, Boucher C, Noyes N, Prosperi M. “The K-mer antibiotic resistance gene variant analyzer (KARGVA),” Front Microbiol., 14:1060891, Mar 2023, doi: 10.3389/fmicb.2023.1060891.

41. K. Břinda, M. Sykulski, and G. Kucherov, “Spaced seeds improve k-mer-based metagenomic classification,” Bioinformatics, vol. 31, no. 22, pp. 3584–3592, Nov. 2015, doi: 10.1093/bioinformatics/btv419.

42. S. Girotto, M. Comin, and C. Pizzi, “Efficient computation of spaced seed hashing with block indexing,” BMC Bioinformatics, vol. 19, no. 15, p. 441, Nov. 2018, doi: 10.1186/s12859-018-2415-8.

